## Supplemental Fig. 1 for "Structural evidence for elastic tethers connecting separating chromosomes in crane-fly spermatocytes"

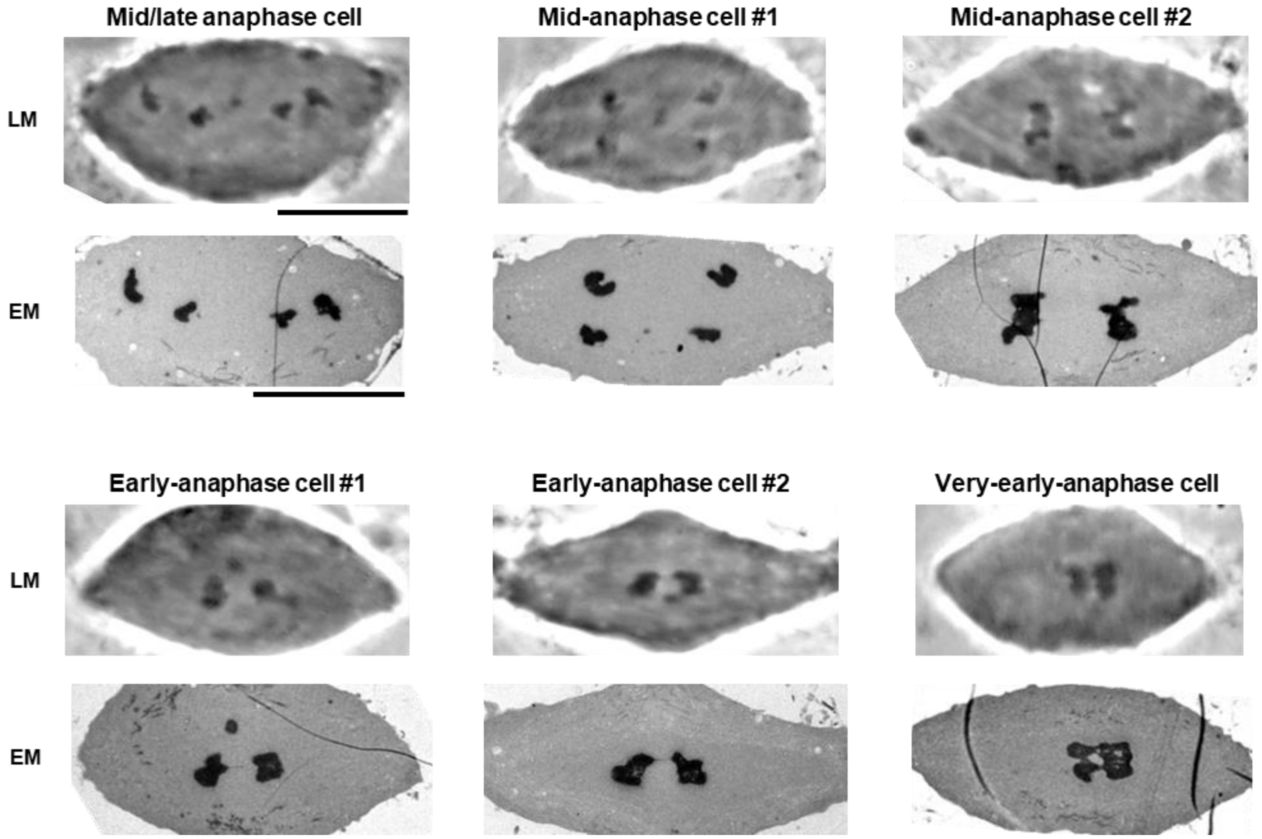


**Supplementary Fig. 1. Correlative live cell imaging with electron microscopy.**
Light and electron microscopy (LM and EM) images of a mid/late-anaphase cell (Fig. 1), two mid-anaphase cells (Figs. 2, 3), two early-anaphase cells (Fig. 4A,C), and a very-early-anaphase cell (Fig. 4E). Scale bars: 10 μm.

**Legend of supplementary Movie 1.**

EM tomographic slices of tethers forming between separating chromosomes in a mid/late-anaphase cell. In the movie, 24-nm thick tomographic section is shown every 0.6 nm. Darker- and lighter-stained filaments are segmented and their 3D meshes are shown in orange and blue, respectively. Scale bar, 200 nm.
